# Polyvinylpyrrolidone enhances AAV transduction *in vitro* and shows preliminary utility for subretinal delivery

**DOI:** 10.64898/2026.07.21.739700

**Authors:** Natalia Gogoleva, Thi-Hang Tran, Mayuko Oki, Shinichi Fukuda, Zeynab Javanfekr Shahri, Eugenia Kumaga, Satoru Takahashi, Michito Hamada

**Affiliations:** Ph.D. Program in Human Biology, Graduate School of Comprehensive Human Sciences, University of Tsukuba, 1-1-1, Tennodai, Tsukuba, Ibaraki, 305-8575, Japan; Department of Anatomy and Embryology, Faculty of Medicine, University of Tsukuba, 1-1-1, Tennodai, Tsukuba, Ibaraki, 305-8575, Japan; Laboratory of Advanced Vision Science, Institute of Medicine, University of Tsukuba, 1-1-1, Tennodai, Tsukuba, Ibaraki, 305-8575, Japan; Laboratory of Regenerative Medicine and Stem Cell Biology, Institute of Medicine, University of Tsukuba, 1-1-1, Tennodai, Tsukuba, Ibaraki, 305-8575, Japan; Laboratory Animal Resource Center, Faculty of Medicine, University of Tsukuba, 1-1-1, Tennodai, Tsukuba, Ibaraki, 305-8575, Japan; International Institute for Integrative Sleep Medicine (WPI-IIIS), University of Tsukuba, 1-1-1, Tennodai, Tsukuba, Ibaraki, 305-8575, Japan; Medical Science Center, Ibaraki Prefectural University of Health Sciences, 4669-2 Ami, Ami-machi, Inashiki-gun, Ibaraki, 300-0394, Japan

**Keywords:** polyvinylpyrrolidone, adeno-associated virus, subretinal injection, synthetic polymer

## Abstract

Adeno-associated virus (AAV) vectors are widely used for gene delivery, but inefficient transduction can require high vector doses. We tested whether unmodified linear polyvinylpyrrolidone (PVP), a pharmaceutical excipient, can improve AAV formulation without chemical modification of the vector or polymer. PVP10, PVP40, and PVP360 were evaluated *in vitro* across HEK293, HeLa, MEF, and CHO cells; selected formulations were tested after intravenous delivery, and 3% PVP40 was tested by subretinal delivery. *In vitro*, 1-3% PVP increased AAV8- and AAV9-mediated GFP expression across multiple cell lines. PVP360 showed broad activity in the initial cross-cell assay, whereas the HEK293 molecular-weight screen identified PVP40 and PVP360 as the most active formulations. The substantial fold increase observed in CHO cells largely reflected the low baseline transduction of the control group. MTT absorbance declined with concentrated PVP360, whereas PVP40 retained transduction-enhancing activity and was selected for local testing. In the HeLa AAV-DJ assay, the response pattern differed between MOI 1,000 and MOI 100; the largest observed increases in GFP-positive area occurred with 1.5% PVP10 and 1.5% PVP360 at MOI 100. Intravenous AAV9 delivery with PVP did not consistently increase *ex vivo* organ reporter signal. By contrast, subretinal delivery of AAV-PHP.eB with 3% PVP40 produced a 1.79- fold larger mean DsRed-positive area per retinal section, averaged within each eye (Welch’s t-test *p* = 0.063; Bayesian Pr[Δ > 0] = 0.952). These findings support further evaluation of PVP40 for local subretinal AAV delivery.

## Introduction

Gene therapy using adeno-associated virus (AAV) vectors has emerged as a leading strategy for the treatment of monogenic disorders and is playing a central role in gene replacement therapies, particularly for autosomal recessive diseases^1^. AAVs are non-enveloped, single-stranded DNA viruses of the Parvoviridae family, valued for their broad tissue tropism, low immunogenicity, and ability to mediate long-term transgene expression from episomal genomes^1,2^. These properties have led to the clinical approval of multiple AAV-based therapies.

Despite these advantages, several critical technical and biological bottlenecks remain for AAV vectors. These include suboptimal or off-target tissue tropism, the frequent requirement for high vector doses, and the associated risks of dose-dependent immunogenicity, hepatotoxicity, and high manufacturing costs^3,4^. To overcome these challenges, extensive efforts have been devoted to improving AAV performance through optimization of production pipelines, capsid engineering, and the use of supplementary agents to enhance or modulate vector transduction^2,5^.

Polyvinylpyrrolidone (PVP) is a synthetic, linear polymer characterized by high biocompatibility, water solubility, and low cytotoxicity^6,7^. Owing to its favorable safety profile, PVP has been widely used as a plasma volume expander, a pharmaceutical excipient and binder, a component of contact lenses, and an additive in cosmetic and industrial products. Importantly, PVP has a long history of clinical and regulatory acceptance^8^. Its formulation-relevant properties, including macromolecular crowding and protein stabilization reported in other contexts, make it a plausible candidate for testing as a viral vector formulation additive^9,10^.

Previous studies have demonstrated that supplementation of AAV vectors with various polymers can enhance transduction efficiency *in vitro* and *in vivo*, often through effects on vector modification and cellular interactions^11–13^. However, most reported approaches rely on chemically modified, less clinically familiar, or complex polymer formulations. In contrast, the potential impact of unmodified, linear PVP on AAV transduction has not been systematically examined.

In this study, we hypothesized that linear PVP, through formulation effects, could enhance AAV transduction efficiency without chemical modification of the vector or polymer. We employed a panel of AAV serotypes- AAV8 and AAV9 as clinically relevant natural serotypes, AAV-DJ as a high-efficiency chimeric benchmark, and AAV-PHP.eB for enhanced *in vivo* transduction- to evaluate PVP effects across multiple *in vitro* systems and to provide initial *in vivo* evidence in both systemic and local delivery models.

## Materials and methods

### Animal Experiments

Double-reporter C57BL/6N-Gt(ROSA)26Sor<tm1(CAG-EGFP/DsRed)Utr>/Rbrc mice were generated as previously described^14^. These mice, hereafter referred to as GRR mice, express EGFP before and DsRed after Cre-mediated recombination. Mice were maintained under specific-pathogen-free conditions at 23.5 ± 2.5 °C and 52.5 ± 12.5% relative humidity with a 14-h light/10-h dark cycle. All procedures complied with national and institutional regulations and were approved by the University of Tsukuba Animal Ethics Committee (authorization no. 25-405).

### AAV Production

AAV vectors were produced using the standard triple-plasmid transfection method. The following rep/cap plasmids were used: pRC-AAV2/8 (Addgene plasmid #112864), pRC-AAV2/9 (Addgene plasmid #112865), and pRC-AAV-DJ (kindly provided by Dr. Yoan Cherasse, IIIS, University of Tsukuba), pRC- AAV-PHP.eB (kindly provided by Dr. Yoan Cherasse, IIIS, University of Tsukuba). The adenoviral helper plasmid (pHelper) and the transgene plasmid pAAV-CMV-GFP were purchased from Takara Bio. The pAAV-CMV-iCre plasmid was generated as previously described^15^. Transfections were performed using PEI MAX (Polysciences, USA) according to the manufacturer’s instructions.

AAV particles were purified using a sucrose-cushion ultracentrifugation method with minor modifications^16^. Briefly, transfected cells were harvested and lysed under high-salt conditions, followed by treatment with TurboNuclease (25 U/mL; Accelagen, USA) for 2 hours at 37 °C to degrade residual nucleic acids. Lysates were clarified by two sequential centrifugation steps at 11,000 × g for 10 min to remove cellular debris while minimizing viral loss. Viral genome titers were determined using an established ITR- targeted qPCR method^17^.

### Polymer Solution Preparation

For *in vitro* experiments, PVP10, PVP40, and PVP360 (Sigma-Aldrich, USA; approximate average molecular weights 10, 40, and 360 kDa, respectively) were dissolved in high-glucose DMEM (Gibco, USA) to generate 10% stock solutions and diluted to working concentrations before use. For *in vivo* experiments, polymers were dissolved in PBS-MK buffer. PBS-MK was prepared by dissolving 26.3 mg MgCl₂ and 14.91 mg KCl in 1× PBS to a final volume of 100 mL, sterilized through a 0.22-µm filter, and stored at 4 °C.

### Cell Culture

HEK293 cells were cultured in high-glucose DMEM supplemented with 10% FBS (BioSera, USA), 2 mM L-glutamine (Gibco, USA), 1 mM MEM-NEAA (Gibco, USA), 1 mM sodium pyruvate (Gibco), and 1× penicillin–streptomycin (Gibco). CHO, primary MEFs, and HeLa cells were maintained in DMEM supplemented with 10% FBS, 2 mM L-glutamine, and 1× penicillin–streptomycin. All cell lines were cultured at 37 °C with 5% CO₂ supply.

### *In vitro* infections, fluorescence imaging, and flow cytometry

For flow-cytometric analysis (FCM), 1-2 × 10^5^ cells were seeded per well in 24-well plates (Gibco). The following day, cells were infected with AAV vectors with or without polymer formulations at the indicated MOIs, incubated at 37 °C for 2 hours, and then supplied with fresh culture medium. For the Figure 1A imaging assay, HEK293 cells received AAV9-CMV-GFP at MOI 40,000 with 0.1%, 1%, or 3% PVP360; the inoculum was replaced after 2 hours and GFP images were acquired at 24 hours post-infection. Images were acquired with a Nikon DS-Fi3 camera through a 10× objective using a manual exposure of 90 ms and a gain of 9.18×; the same settings were used for all conditions. For the Figure 1B-E flow-cytometry assays, MOI 40,000 was used for AAV9-infected HEK293 cells, AAV8-infected HEK293 cells, and AAV9-infected primary MEFs, and MOI 20,000 was used for AAV9-infected CHO cells. Three recorded values are shown per condition, except that the Figure 1C 1% PVP360 condition contained two recorded values. For the Figure 2B HEK293 molecular-weight screen, AAV9-CMV-GFP was used at MOI 10,000 with 1% or 3% PVP10, PVP40, or PVP360, and cells were analyzed at 3 days post-infection. For the HeLa flow-cytometry experiment, AAV8-CMV-GFP- and AAV9-CMV-GFP-infected cells were analyzed at 3 days post-infection, with three recorded values per condition (Supplementary Figure 3). For the continuous co-administration experiment (Supplementary Figure 2), AAV9 at MOI 10,000 was resuspended in a 20 µL PBS formulation containing no polymer, 3% PVP40, or 3% PVP360, incubated on ice for 1 hour, and added directly to plated cells without subsequent medium replacement. The stated 3% PVP concentration refers to the 20 µL premix before addition to the well. Cells were analyzed by flow cytometry at 4 days post-infection.

**Figure 1.**
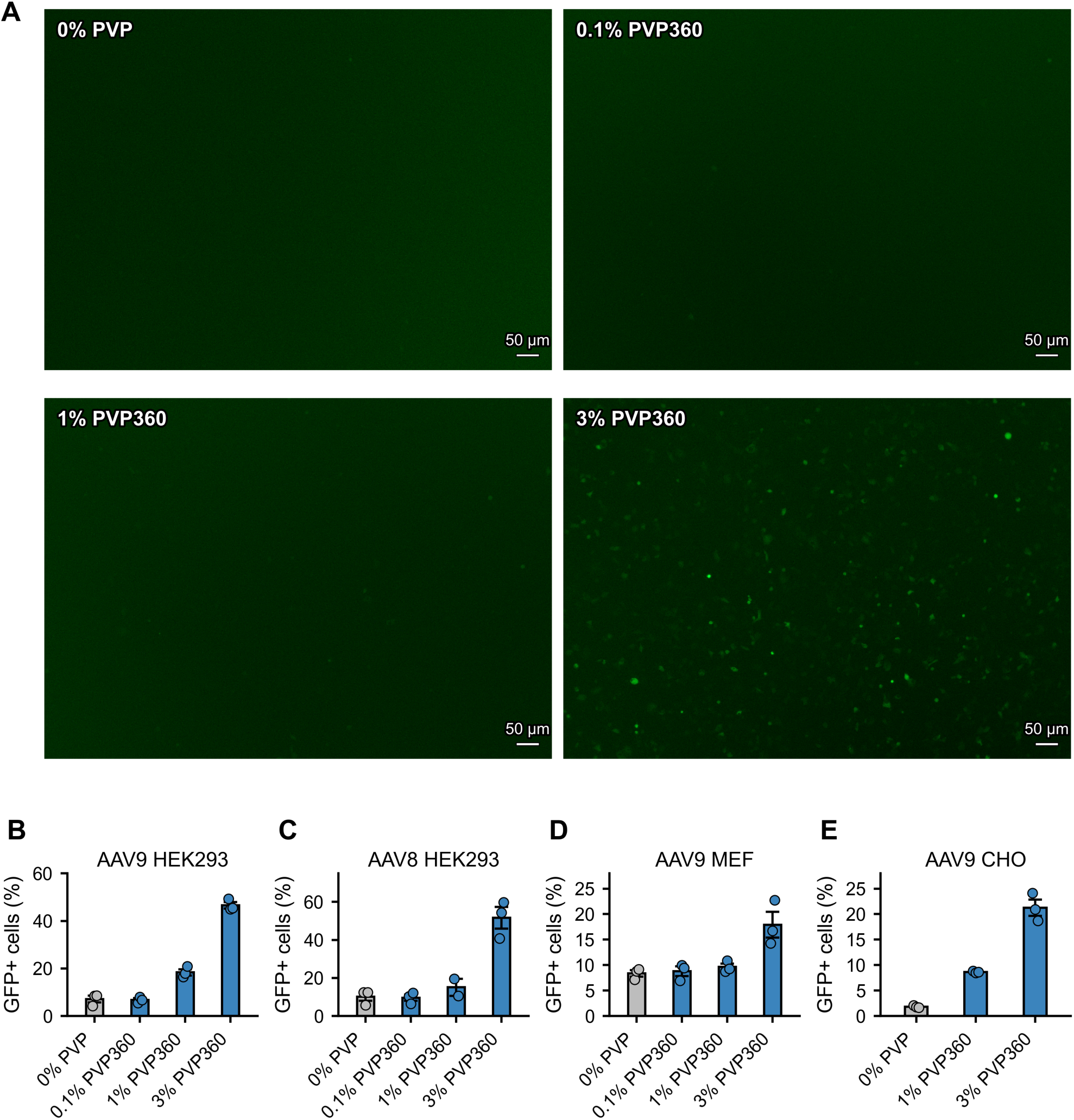
PVP360 increases *in vitro* AAV reporter transduction. (A) Representative AAV9-CMV-GFP fluorescence images acquired at 24 hours post-infection with 0%, 0.1%, 1%, or 3% PVP360 using identical camera settings; scale bars, 50 µm. (B-E) Flow-cytometric quantification of GFP-positive cells for AAV9-infected HEK293 cells, AAV8-infected HEK293 cells, AAV9-infected MEFs, and AAV9-infected CHO cells. MOI was 40,000 for panels A-D and 20,000 for panel E. The 0% PVP groups contain AAV without polymer. Bars show mean ± SEM with individual values.

**Figure 2.**
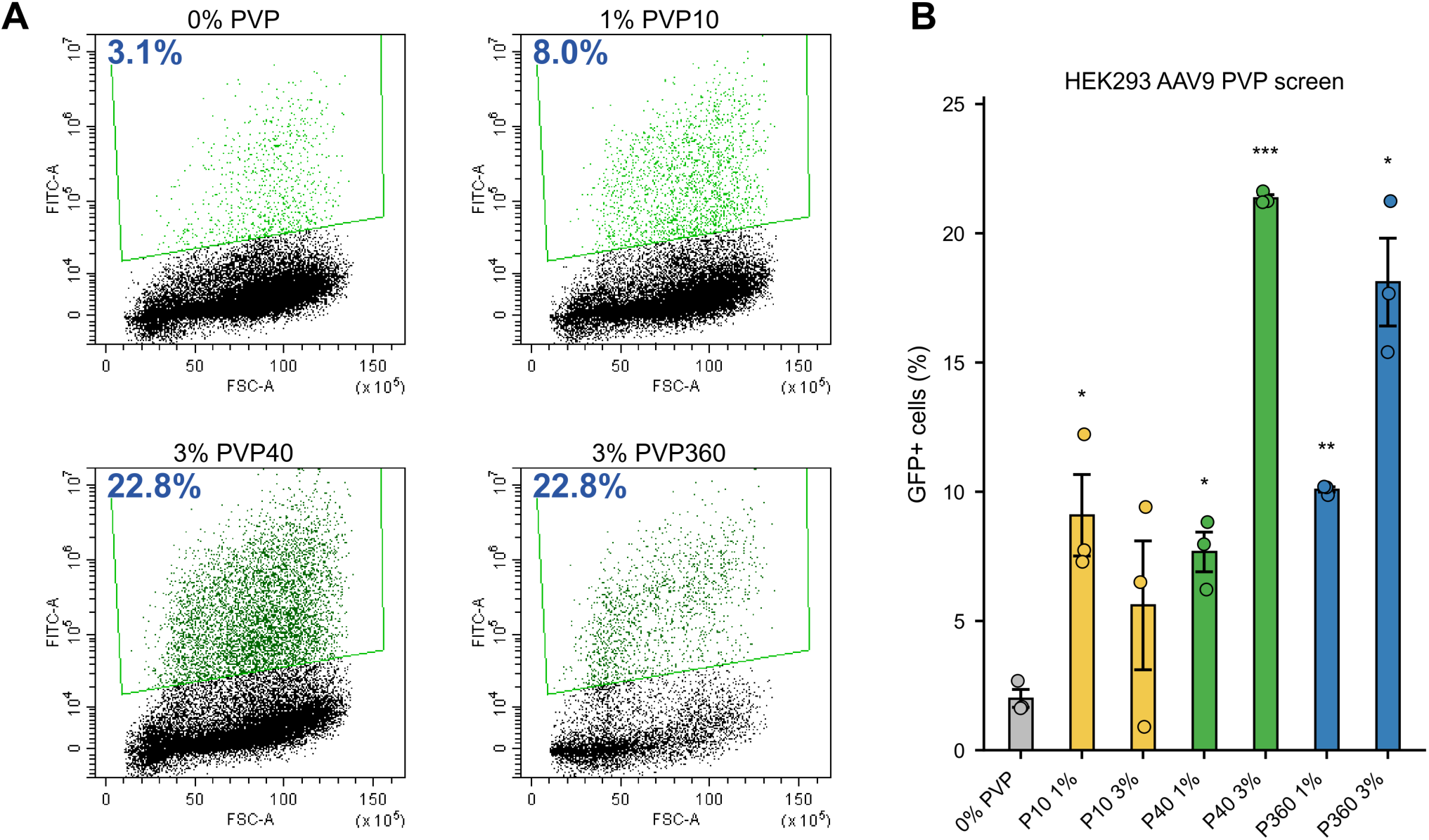
PVP molecular-weight screen in HEK293 cells. (A) Representative final GFP gates for AAV9-infected HEK293 cells; values indicate the percentage of events within the GFP-positive gate. (B) GFP-positive HEK293 cells across PVP10, PVP40, and PVP360 concentrations. The 0% PVP group contains AAV9 without polymer. Asterisks in panel B indicate BH-adjusted *q* values from Welch’s t-tests versus 0% PVP (* *q* < 0.05, ** *q* < 0.01, *** *q* < 0.001). HeLa FCM data and the MTT/viable-cell data are shown in Supplementary Figures 3 and 4.

At the indicated time point (3 or 4 days post-infection), cells were detached using a cell scraper, gently resuspended by pipetting, transferred to 5 mL tubes through a 40-µm filter, washed with flow-cytometry buffer (DPBS containing 1% FBS; BioSera, USA), and resuspended in flow-cytometry buffer containing 1 µg/mL DAPI (Wako, Japan). Data were acquired on a CytoFLEX cytometer (Beckman Coulter, USA) and analyzed using CytExpert software. At least 4 × 10^4^ events were acquired per sample. The analysis sequence comprised the main cell population, singlets, DAPI-negative viable cells, and the final GFP-positive gate. The viable-cell data in Supplementary Figure 4 are reported as the percentage of DAPI-negative cells. Representative final GFP gates supporting Figure 1 are shown in Supplementary Figure 1, and representative HEK293 gates for the molecular-weight screen are shown in Figure 2A.

### *In vitro* infectivity assay of HeLa cells

HeLa cells were plated in 96-well Nunclon Delta-treated flat-bottom plates (ThermoFisher Scientific, USA) at 5 × 10^4^ cells per well. Cells were infected with AAV-DJ-CMV-GFP at an MOI of 1,000 (standard condition) or 100 (10-fold lower condition) in the presence or absence of PVP10, PVP40, or PVP360 at final concentrations of 0.25%, 0.5%, or 1.5%, without a medium change. GFP fluorescence was imaged at 72 hours post-infection using a BZ-X800 microscope (Keyence, Japan) with matched acquisition settings. Three images were quantified per condition. GFP-positive area was quantified using the BZ-X800 Analyzer software with identical threshold settings across groups.

### MTT assay

Cells were plated at 5 × 10^4^ cells per well in 96-well Nunclon Delta-treated flat-bottom plates (Thermo Fisher Scientific, USA). PVP10, PVP40, or PVP360 was tested at final concentrations of 0.5%, 1.5%, 2.5%, 4%, or 5%, with parallel groups receiving AAV9-CMV-GFP. Cells were incubated overnight, MTT reagent was applied according to the manufacturer’s instructions (Nacalai Tesque, Japan), and absorbance was measured at 540 nm. HEK293 groups contained three recorded values per condition. HeLa groups contained three recorded values except the 0% PVP + AAV and 0.5% PVP40 + AAV conditions, which contained two values.

### Intravenous Injections

Eight-week-old male mice were pre-warmed using a hot-water bag to promote vasodilation. Tail-vein injections were performed in a total volume of 200 µL per mouse, containing 5 × 10^9^ vg AAV9-CMV-iCre in PBS-MK alone or with 3% PVP40 or 3% PVP360. Vector-polymer formulations were pre-incubated on ice for 2 hours. PVP40 and PVP360 were evaluated in separate mouse cohorts with separate PBS-MK control groups. The PVP40 dataset comprised 4 control and 5 treatment mice for each organ; the PVP360 dataset comprised 5 control and 5 treatment mice for each organ except brain, for which 4 control and 5 treatment measurements were available. All animals were monitored on the day after injection.

### Subretinal Injections

General anesthesia in 10-12-week-old male mice was induced by intraperitoneal administration of midazolam (4 mg/kg; SANDOZ), medetomidine hydrochloride (0.3 mg/kg; Nippon Zenyaku Kogyo), and butorphanol tartrate (5 mg/kg; Meiji Seika Pharma). Atipamezole hydrochloride (0.3 mg/kg) was administered for recovery. Subretinal injections were performed following previously established procedures^18^. Each eye received 1 µL containing 7 × 10^7^ vg AAV-PHP.eB-CMV-iCre in PBS-MK vehicle or with 3% PVP40 after pre-incubation on ice for 2 hours. Both eyes of each mouse received the same formulation; each group comprised 3 mice and 6 eyes. Mice were randomly allocated to formulation groups, and retinal outcome assessment was performed with group identity masked. No animals, eyes, or retinal sections were excluded from the analysis. All animals were monitored on the day after injection, and eyes were collected 2 weeks after injection.

### *Ex vivo* organ imaging after intravenous injection

One month after tail-vein injection, mice were euthanized and perfused with PBS to remove circulating blood. Organs were harvested, and DsRed fluorescence was measured using the NEWTON 7.0 imaging system. Liver fluorescence was acquired at 620 nm excitation, whereas brain, lung, kidney, gastrointestinal tract (GI), and heart fluorescence were acquired at 580 nm. Signal quantification and background subtraction were performed using Quant software. PVP40 data were reported as background-subtracted radiance and PVP360 data as background-subtracted total counts; the two acquisition datasets were not compared directly across formulations.

### Cryosection Preparation and Imaging

Eyes were fixed in 4% paraformaldehyde (PFA) in PBS for 4 h at 4 °C, followed by sequential cryoprotection in 20% sucrose/PBS overnight, then 30% sucrose/PBS overnight at 4 °C. Tissues were embedded in OCT compound, and cryosections (10 µm) were prepared. Sections were briefly dried, rinsed with PBS, and mounted in Vectashield mounting medium (Vector Laboratories, USA). Images were acquired using a BZ-X800 microscope. For each eye, DsRed-positive area was measured in seven retinal sections distributed across seven slides using the same Keyence condition file and identical analysis settings across groups. The seven values were averaged to calculate the mean DsRed-positive area per retinal section for that eye. Eye sections were stained using a standard hematoxylin and eosin (H&E) protocol. For total retinal thickness, three retinal positions were measured per eye and averaged to produce one eye-level value (n = 6 eyes/group).

### Statistical Analysis

All quantitative data are presented as mean ± standard error of the mean (SEM) unless otherwise stated. For the Figure 2B molecular-weight screen, Welch’s t-tests were performed versus the 0% PVP control, and *p*-values were adjusted across the six comparisons using the Benjamini-Hochberg method. Asterisks in Figure 2B indicate BH-adjusted *q* values (* *q* < 0.05, ** *q* < 0.01, *** *q* < 0.001). For the Figure 3 screening panels, Welch *p* values and BH-adjusted *q* values are provided in Supplementary Table S1. For the subretinal experiment, AAV+PBS-MK and AAV+PVP40 groups were compared using Welch’s t-test with n = 6 eyes/group; a Mann-Whitney test was also performed.

**Figure 3.**
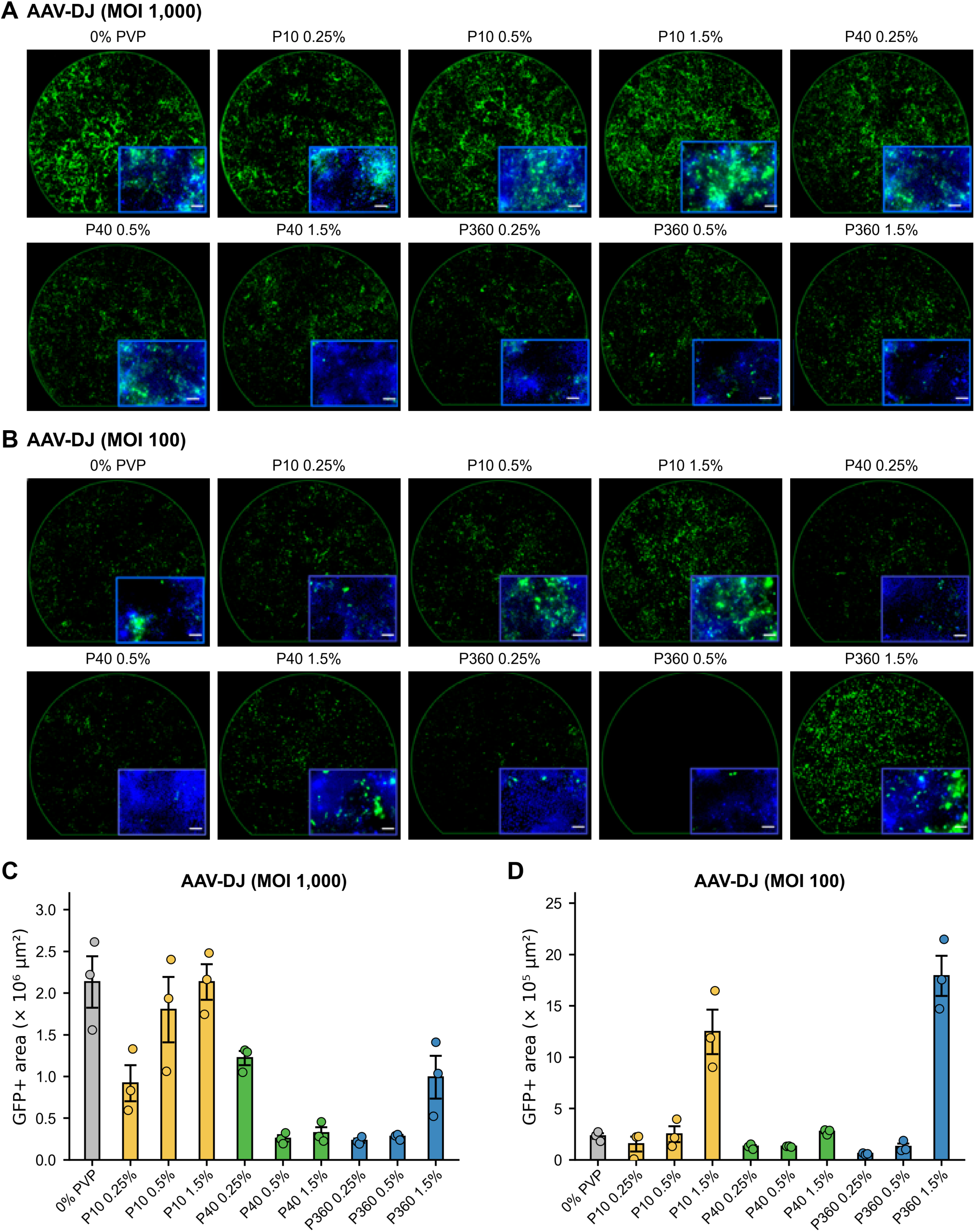
AAV-DJ infectivity assay. (A-B) Representative fluorescence images at MOI 1,000 and MOI 100, respectively. Main fields show GFP fluorescence. Insets show GFP with DAPI nuclear counterstaining and were acquired at 40×; inset scale bars, 50 µm. The 0% PVP condition contains AAV-DJ without polymer. (C-D) GFP-positive area at MOI 1,000 and MOI 100. Bars show mean ± SEM with three recorded image measurements per condition. Welch *p* values and Benjamini-Hochberg *q* values are provided in Supplementary Table S1; no Figure 3 comparison remained significant after BH adjustment.

For in vivo datasets, comparisons were analyzed according to the replicate unit defined for each experiment. The primary subretinal endpoint was the mean DsRed-positive area per retinal section, calculated for each eye. Bayesian sensitivity analyses were performed for in vivo comparisons. Observed values were modeled independently by group as ygi ∼ Normal(μg, σg2), using the reference prior p(μg, σg2) ∝ 1/σg2. Treatment effects were defined as Δ = μtreatment - μcontrol. The marginal posterior for each group mean was sampled as μg = mean(yg) + [sg/sqrt(ng)] × t, where t follows a Student t distribution with ng - 1 degrees of freedom and sg is the sample standard deviation. For the intravenous comparisons, 400,000 Monte Carlo draws per group were generated; for the subretinal comparison, 600,000 draws per group were generated. All Bayesian simulations used random seed 20260708. Posterior probability Pr(Δ > 0), the median Δ, and central 89% and 95% credible intervals were calculated directly from posterior Δ samples. Kernel density estimation was used only to draw posterior density curves. Background-subtracted IV values, including negative values, were retained on the original measurement scale. Bayesian analyses were implemented in Python 3.12.10 using NumPy 2.3.5, pandas 2.3.3, SciPy 1.16.3, Matplotlib 3.10.8, and Pillow 12.0.0.

## Results

### PVP enhances AAV transduction across serotypes and cell types

To determine whether unmodified PVP could increase AAV transduction, we first tested PVP360 with AAV9-CMV-GFP in HEK293 cells. After a 2-hour exposure at MOI 40,000 followed by medium replacement, GFP expression increased with PVP360 concentration and was readily detectable with 3% PVP360 at 24 hours post-infection (Figure 1A). To test the breadth of this effect, we compared AAV capsids and cell types. At 3 days post-infection, 1% and 3% PVP360 increased the proportion of GFP-positive HEK293 cells by 2.5-fold and 6.5-fold for AAV9 and by 1.5-fold and 5.0-fold for AAV8, respectively (Figure 1B-C). PVP360 also increased AAV9 transduction in primary MEFs and CHO cells; in CHO cells, the substantial fold increase largely reflected the low baseline transduction of the 0% PVP control (Figure 1D-E). Representative final GFP gates are shown in Supplementary Figure 1.

Because the initial assays used medium replacement after a 2-hour exposure, we also tested continuous co-administration. Without subsequent medium replacement, the mean GFP-positive fraction was 1.8-fold higher with the PVP40 premix and 2.0-fold higher with the PVP360 premix than with the 0% PVP premix (Supplementary Figure 2).

### Formulation screening supports PVP40 selection for local testing

These results prompted a direct comparison of PVP10, PVP40, and PVP360 across concentrations to identify a formulation for *in vivo* testing. In the HEK293 AAV9 screen, PVP40 and PVP360 produced the largest increases in GFP-positive cells, whereas PVP10 showed smaller and more variable effects (Figure 2A-B). The HeLa flow-cytometry experiment showed a similar pattern for AAV8 and AAV9 under selected conditions (Supplementary Figure 3). We next compared the *in vitro* metabolic-activity profiles of the three polymers. MTT absorbance declined at the upper PVP360 concentrations but was better preserved with PVP40. The DAPI-negative fraction was similar across PVP conditions in HeLa cells and more variable in HEK293 cells (Supplementary Figure 4).

We then asked whether the effect extended to a different capsid across two vector inputs. In an AAV- DJ^19^ HeLa screen, the response pattern differed between MOI 1,000 and MOI 100. At MOI 1,000, PVP40 and PVP360 generally reduced GFP-positive area relative to the 0% PVP control, whereas at MOI 100 the largest increases occurred with 1.5% PVP10 and 1.5% PVP360 (Figure 3). None of the individual comparisons remained significant after Benjamini-Hochberg adjustment; Welch *p* values and BH-adjusted *q* values for Figures 2 and 3 are provided in Supplementary Table S1. Overall, PVP activity varied across assay conditions. PVP40 was selected for local *in vivo* testing because it enhanced AAV8 and AAV9 transduction while preserving MTT absorbance better than PVP360.

### PVP40 increases mean retinal reporter signal after subretinal delivery

The *in vitro* formulation screen identified PVP40 for local *in vivo* testing. AAV-PHP.eB-CMV-iCre, an engineered capsid that efficiently transduces mouse retina^20^, was formulated in PBS-MK vehicle or with 3% PVP40 and delivered by subretinal injection. At two weeks, AAV+PVP40 increased the mean DsRed-positive area per retinal section by 1.79-fold relative to AAV+PBS-MK vehicle (Figure 4). The Welch comparison yielded *p* = 0.063, while Bayesian analysis favored a positive group mean difference (Pr[Δ > 0] = 0.952; Figure 4B and Supplementary Figure 7). The effect size and posterior distribution suggest a positive effect of PVP40 on local retinal reporter expression. Retinal morphology and total thickness were comparable between groups at two weeks (Supplementary Figure 8).

**Figure 4.**
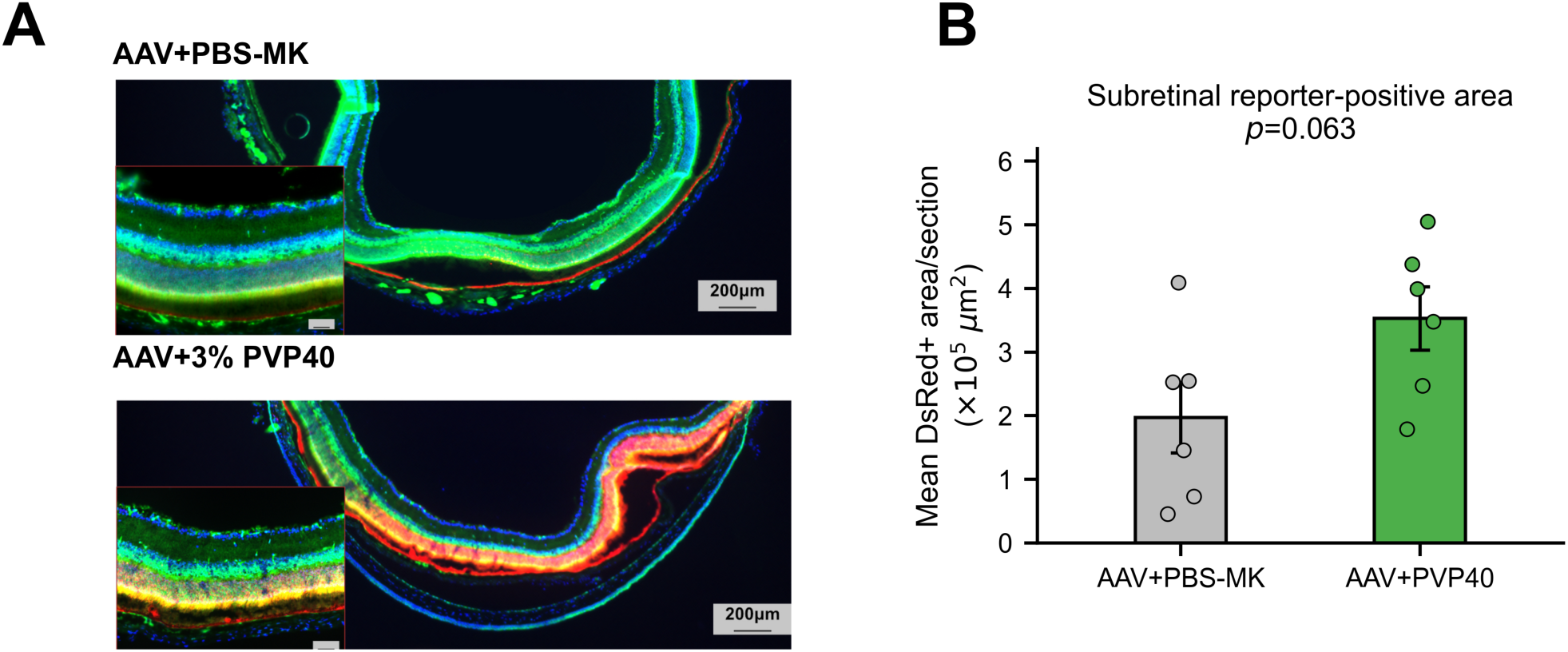
Local subretinal delivery with 3% PVP40. (A) Representative low-magnification and inset retinal fluorescence images acquired and displayed with matched settings. Green denotes baseline EGFP, red denotes Cre-dependent DsRed, and blue denotes nuclear counterstaining. Scale bars, 200 µm (overview) and 50 µm (inset). (B) Mean DsRed-positive area per retinal section, averaged within each eye (n = 6 eyes/group). AAV+PVP40 produced a 1.79-fold larger mean than AAV+PBS-MK vehicle; Welch’s t-test *p* = 0.063 and Mann-Whitney *p* = 0.132. Bayesian analysis is shown in Supplementary Figure 7. Data are mean ± SEM with individual eye-level values shown.

To determine whether PVP enhancement also occurred after systemic delivery, we tested PVP40 and PVP360 with tail-vein AAV9 administration in separate experiments. Neither formulation consistently increased *ex vivo* organ reporter signal. PVP360 produced an isolated positive posterior signal in lung, while the remaining organ comparisons showed no consistent enhancement (Supplementary Figure 5). Representative organ images from the same experiments are shown in Supplementary Figure 6. Thus, PVP40 produced a larger retinal reporter signal in the subretinal experiment, whereas systemic administration did not yield consistent organ-level gains in these experiments.

## Discussion

In this study, supplementation of AAV vectors with unmodified, linear PVP enhanced transduction in multiple *in vitro* systems and was associated with a larger mean reporter-positive area after subretinal delivery. The magnitude and direction of the responses differed across experiments involving different cell types, capsids, PVP molecular weights and concentrations, vector inputs, and administration routes. Unmodified linear PVP enhanced AAV transduction without chemical modification of the vector, and PVP40 was prioritized for subretinal evaluation based on its *in vitro* activity, MTT profile, and retinal reporter result. *In vitro* responses varied across experiments, and neither PVP formulation consistently increased systemic organ reporter signal.

*In vitro*, PVP increased AAV8- and AAV9-mediated transduction across multiple cell lines (Figure 1; Supplementary Figures 1–3). The largest fold increase occurred with 3% PVP360 in CHO cells, where baseline transduction was low, while the HEK293 molecular-weight screen showed strong activity with both PVP40 and PVP360 (Figure 2). This pattern is consistent with reports that other soluble macromolecules can alter AAV transduction. In Yang et al., preincubation of AAV6 with polyvinyl alcohol (PVA) increased transduction, cellular uptake, and nuclear accumulation of vector genomes in hematopoietic cells; Wang et al. showed that human serum albumin bound AAV8 virions and increased AAV8 binding to Huh7 cells and transduction^21,22^. Zou et al. reported that 3.6% PEG4000 increased rAAV9 reporter expression in HeLa, HEK293T, K562, and PC12 cells^13^. Although these additives differ chemically and were tested with different capsids and cell systems, they provide precedent that noncovalent formulation additives can increase AAV transduction without capsid engineering.

Across the separate transduction and MTT assays, higher PVP molecular weight did not uniformly provide the best combination of reporter transduction and preserved MTT absorbance (Figure 2 and Supplementary Figure 4). PVP360 produced strong activity in several assays but reduced MTT absorbance at upper concentrations, whereas PVP40 retained activity with better-preserved MTT absorbance. In the PEG-rAAV9 study, PEG2000, PEG4000, and PEG10000 enhanced transduction in HeLa cells, while CCK- 8 results worsened with increasing PEG molecular weight and concentration^13^. Lee et al. found that soluble, unconjugated PEG did not alter AAV2 transduction in control experiments, whereas covalent PEGylation preserved infectivity up to a critical PEG:lysine conjugation ratio and reduced infectivity above that threshold, potentially through steric interference or modification of capsid lysines involved in infection^23^. Although covalent PEGylation is not directly comparable with free PVP supplementation, these findings show that polymer size, amount, and mode of capsid association can determine whether AAV activity is retained or impaired. Thus, polymer selection should balance transduction gain with preservation of MTT absorbance rather than maximize molecular weight or concentration alone.

In the AAV-DJ assay, the response pattern differed between MOI 1,000 and MOI 100 (Figure 3). Larger numerical increases in GFP-positive area occurred primarily with 1.5% PVP10 and 1.5% PVP360 at MOI 100, whereas most PVP40 and PVP360 conditions reduced GFP-positive area at MOI 1,000; none of the individual comparisons remained significant after Benjamini-Hochberg adjustment (Supplementary Table S1). The response pattern differed by vector input, although this non-factorial experiment did not test an interaction; Zou et al. showed that 40% PEG4000 enhanced low-titer rAAV9 expression after stereotaxic intraparenchymal brain injection, a substantially different formulation and route^13^. A factorial experiment spanning AAV dose and PVP concentration should test this interaction. Because PVP has been used as a macromolecular crowder in non-viral cell-culture systems, excluded-volume effects are one hypothesis for the condition-dependent changes observed here^9^.

The two *in vivo* experiments produced different patterns, but they used different capsids, target tissues, vector doses, and readouts and therefore do not directly compare administration routes. Tail-vein AAV9 delivery with PVP40 or PVP360 did not produce broad organ-level enhancement; however, Bayesian analysis showed a positive posterior shift for PVP360 in lung, indicating a possible organ-selective response (Supplementary Figure 5). Representative *ex vivo* organ images are shown in Supplementary Figure 6. Systemic enhancement has been reported after pre-injection of charged polymers before AAV2 administration and after preincubation of AAV6 with PVA^11,21^. However, in a systemic AAV9 study, Chai et al. found that human serum albumin enhanced liver transduction without significantly increasing whole-body transduction, whereas selected serum proteins and cryoprecipitate increased whole-body AAV9 reporter signal^24^. These studies show that systemic formulation effects can differ by additive, capsid, and organ, consistent with the absence of a uniform systemic PVP response in the present study. In the separate eye-level subretinal analysis, the PVP40 group had a 1.79-fold larger mean DsRed-positive area per retinal section (Figure 4B; Welch’s t-test *p* = 0.063). Bayesian analysis favored a positive group mean difference (Pr[Δ > 0] = 0.952), although the central 95% credible interval included zero (Supplementary Figure 7). In a distinct mouse-brain model, 40% PEG4000 increased rAAV9 transduction efficiency within a smaller measured transduction area after intraparenchymal injection; increased viscosity was proposed, but not measured, as one possible contributor^13^. The present data support confirmatory subretinal PVP40 studies.

Device-associated vector loss is a relevant formulation variable for small-volume subretinal delivery. Reichel et al. reported cannula-associated reductions in recovered vector genomes for clinically prepared AAV2- and AAV8-based products that contained poloxamer/PF68; infectivity was not measured^25^. Earlier voretigene development work showed that addition of 0.001% PF68 restored near-complete AAV2-RPE65 vector-genome recovery after passage through administration devices^26^. These studies show that formulation can affect post-device vector recovery. Whether PVP alters device-associated recovery or post-injection dispersion was not tested here. Because PVP solution viscosity depends on molecular weight and concentration^27^, the actual 3% PVP40/PBS-MK formulation should be characterized rheologically. Quantifying vector genomes and infectivity before and after device passage would test recovery, while separate capsid-binding and uptake assays would test cellular mechanisms.

The *in vitro* experiments were designed as formulation screens across multiple cell types, capsids, vector inputs, and PVP conditions rather than as a mechanistic study. The systemic and subretinal experiments used different capsids, doses, tissues, and reporter readouts and therefore cannot be compared directly. The subretinal study tested one PVP40 concentration with one capsid at a single time point and measured mean DsRed-positive area per section in six eyes from three mice per group; the eye-level Welch and Bayesian analyses did not model within-mouse correlation. This area-based endpoint cannot distinguish an increase in transduced-cell number from changes in per-cell reporter intensity, vector recovery, or post-injection distribution. Taken together, the data nominate PVP40 for larger subretinal studies with increased animal-level replication, clustered analysis, and functional, inflammatory, and longer-term safety endpoints.

## Data Availability

Numerical values underlying the quantitative figure panels and the statistical summaries reported in the manuscript are provided in the accompanying Source Data file. Raw images and instrument exports retained for this study are available from the corresponding authors upon reasonable request.

## Author Contributions

M.H. and S.T. conceived and designed the project. N.G. designed the experiments, produced the AAV vectors, and performed the in vitro transduction and tolerability experiments. M.O. performed selected in vitro experiments and assisted with flow cytometry. M.H. performed the intravenous AAV administration experiments. T.H.T. and S.F. performed the subretinal injection experiments. N.G. performed the ex vivo organ and retinal imaging and the associated quantitative analyses. Z.J.S. maintained and bred the GRR mouse colony, provided animals, and assisted with the mouse experiments. N.G. and M.H. wrote the original manuscript draft. Z.J.S. and E.K. contributed to manuscript editing. M.H. and S.T. contributed to the discussion and overall project direction. All authors reviewed, edited, and approved the final manuscript.

---

**Supplementary Figure 1.**
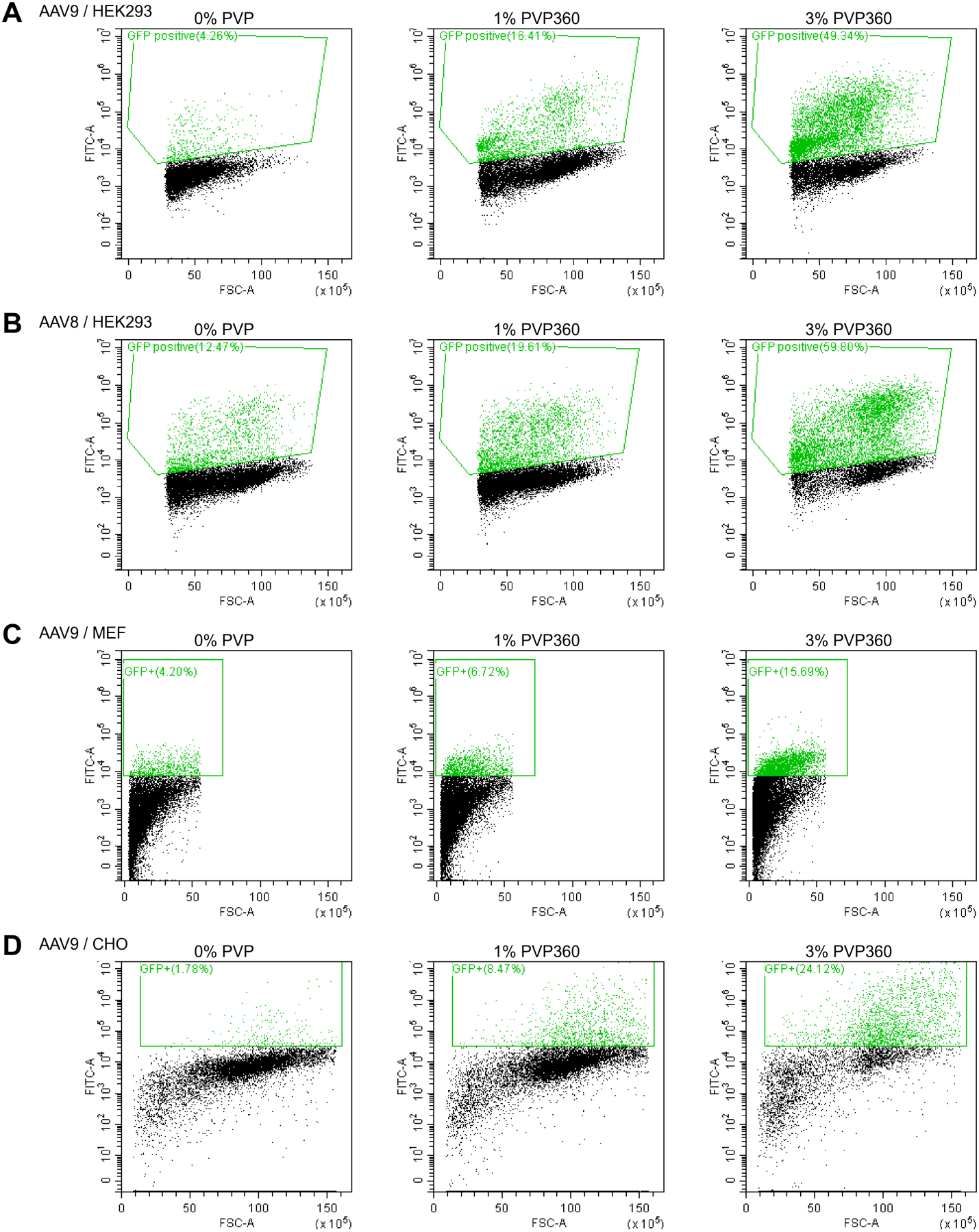
Representative final GFP gates supporting Figure 1. Final GFP gates are shown for AAV9- infected HEK293 cells, AAV8-infected HEK293 cells, AAV9-infected primary MEFs, and AAV9-infected CHO cells. All conditions contain AAV; 0% PVP denotes AAV without polymer.

**Supplementary Figure 2.**
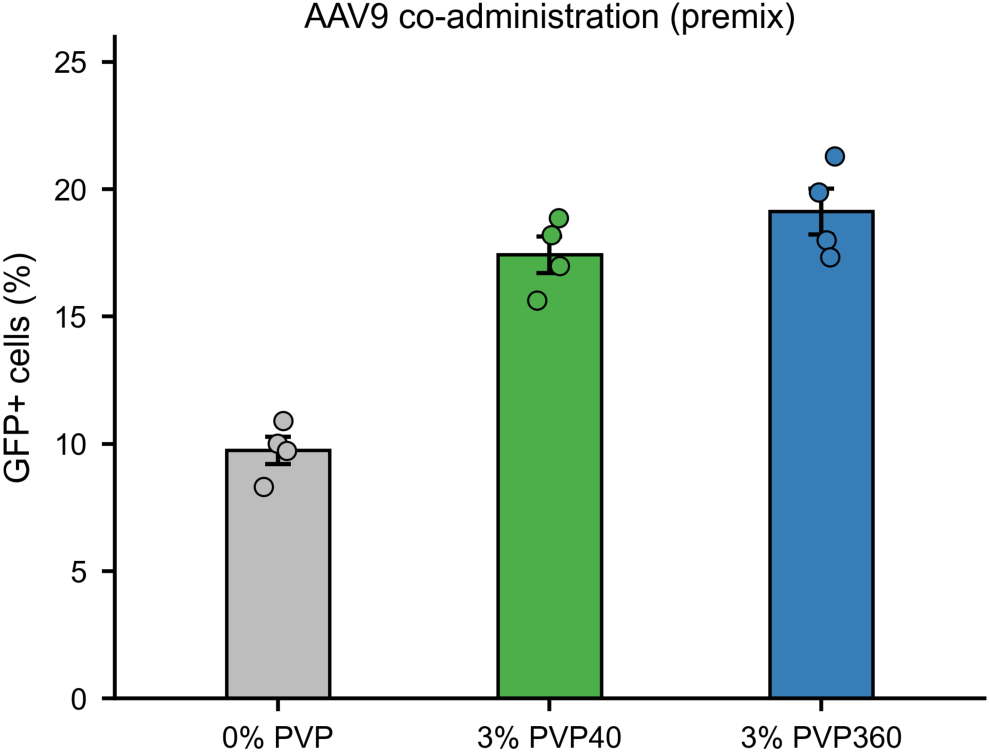
Continuous co-administration without medium replacement. AAV9 was co-administered with 0% PVP, 3% PVP40, or 3% PVP360 and retained without subsequent medium replacement. GFP-positive cells were measured at 4 days post-infection. Bars show mean ± SEM; dots show four plotted replicate samples.

**Supplementary Figure 3.**
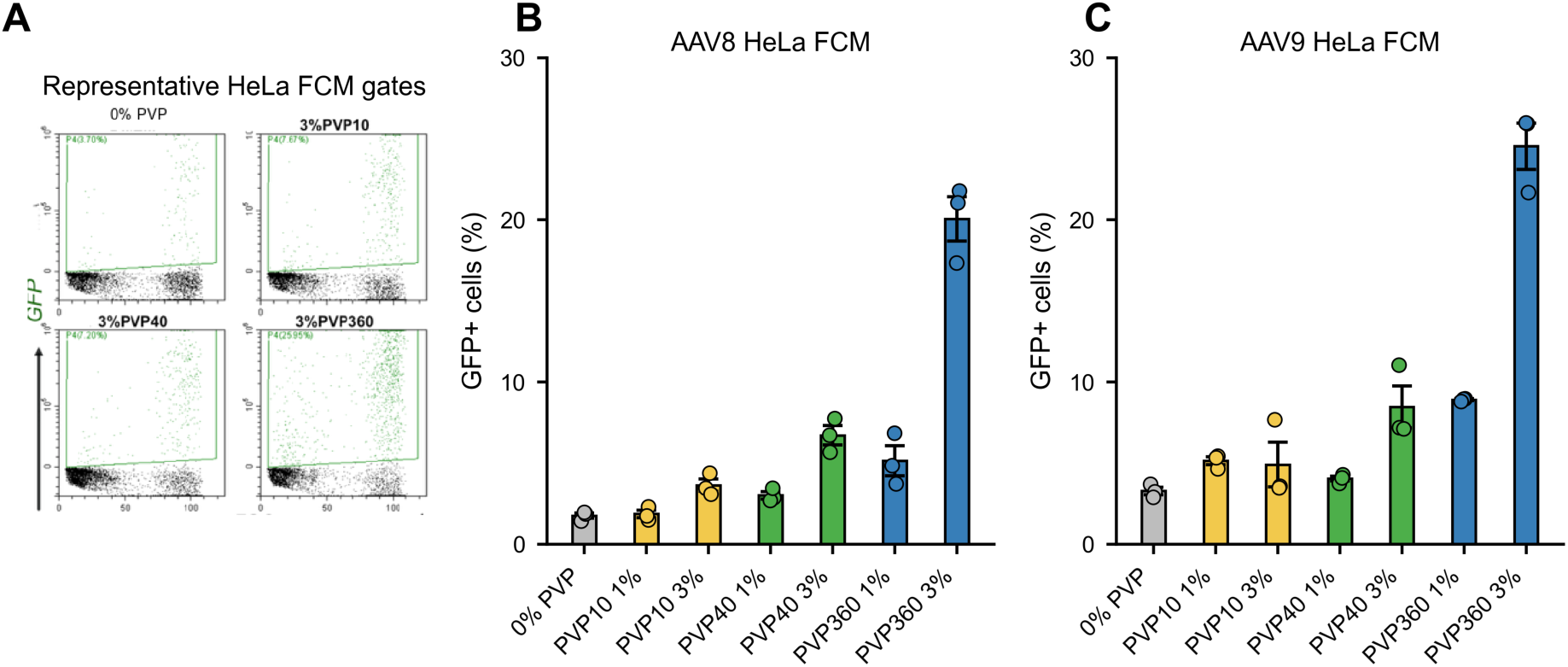
AAV8 and AAV9 transduction in HeLa cells. (A) Representative final GFP gates from the HeLa flow-cytometry experiment. (B-C) GFP-positive HeLa cells after AAV8 or AAV9 infection across PVP10, PVP40, and PVP360 concentrations. The 0% PVP condition contains AAV without polymer. Bars show mean ± SEM with three recorded values per condition.

**Supplementary Figure 4.**
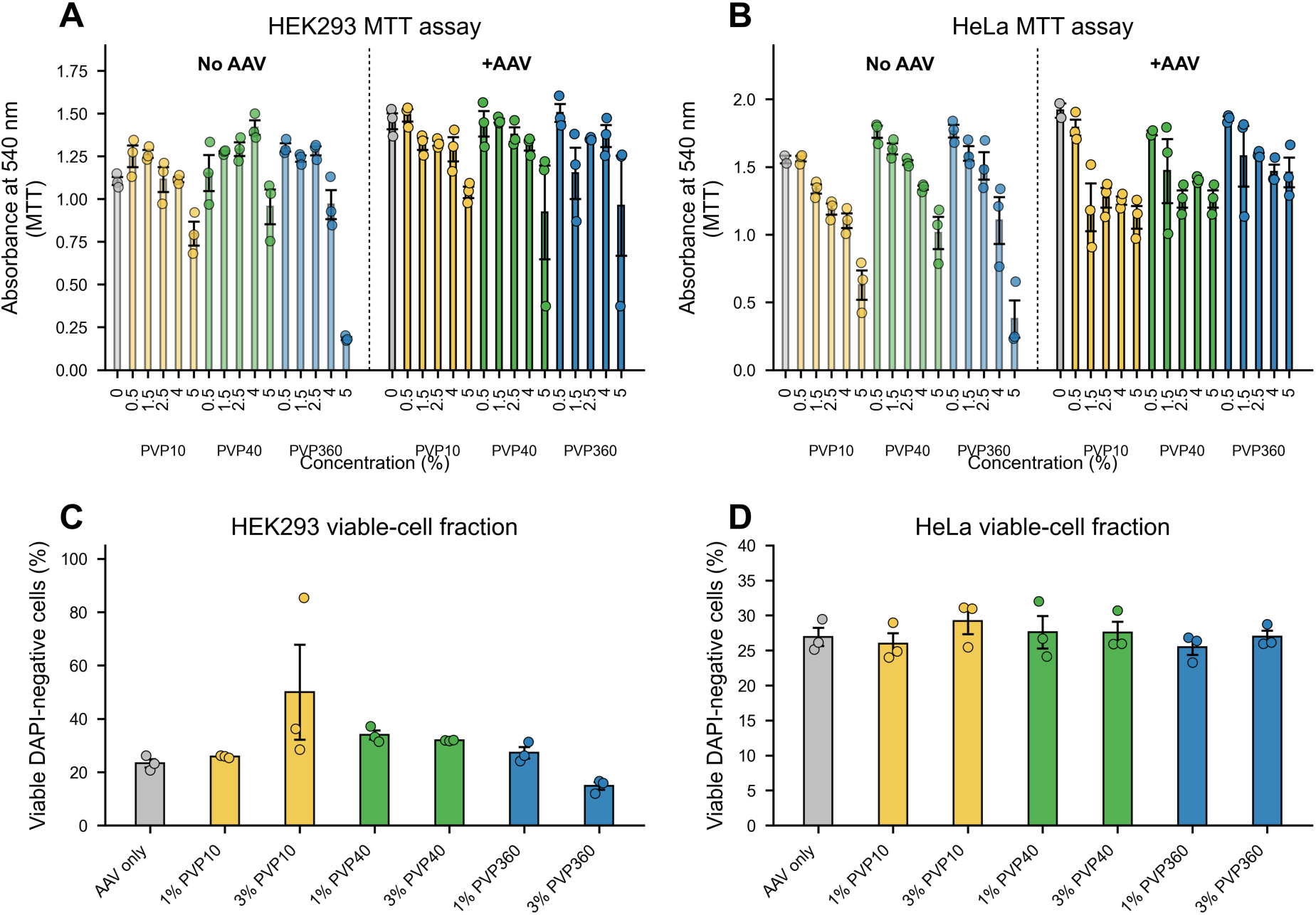
MTT metabolic activity and viable DAPI-negative cell percentages. (A) HEK293 MTT absorbance at 540 nm across PVP concentrations with and without AAV9. (B) HeLa MTT absorbance at 540 nm across PVP concentrations with and without AAV9. (C-D) Percentages of viable DAPI-negative HEK293 and HeLa cells. Bars show mean ± SEM with individual recorded values.

**Supplementary Figure 5.**
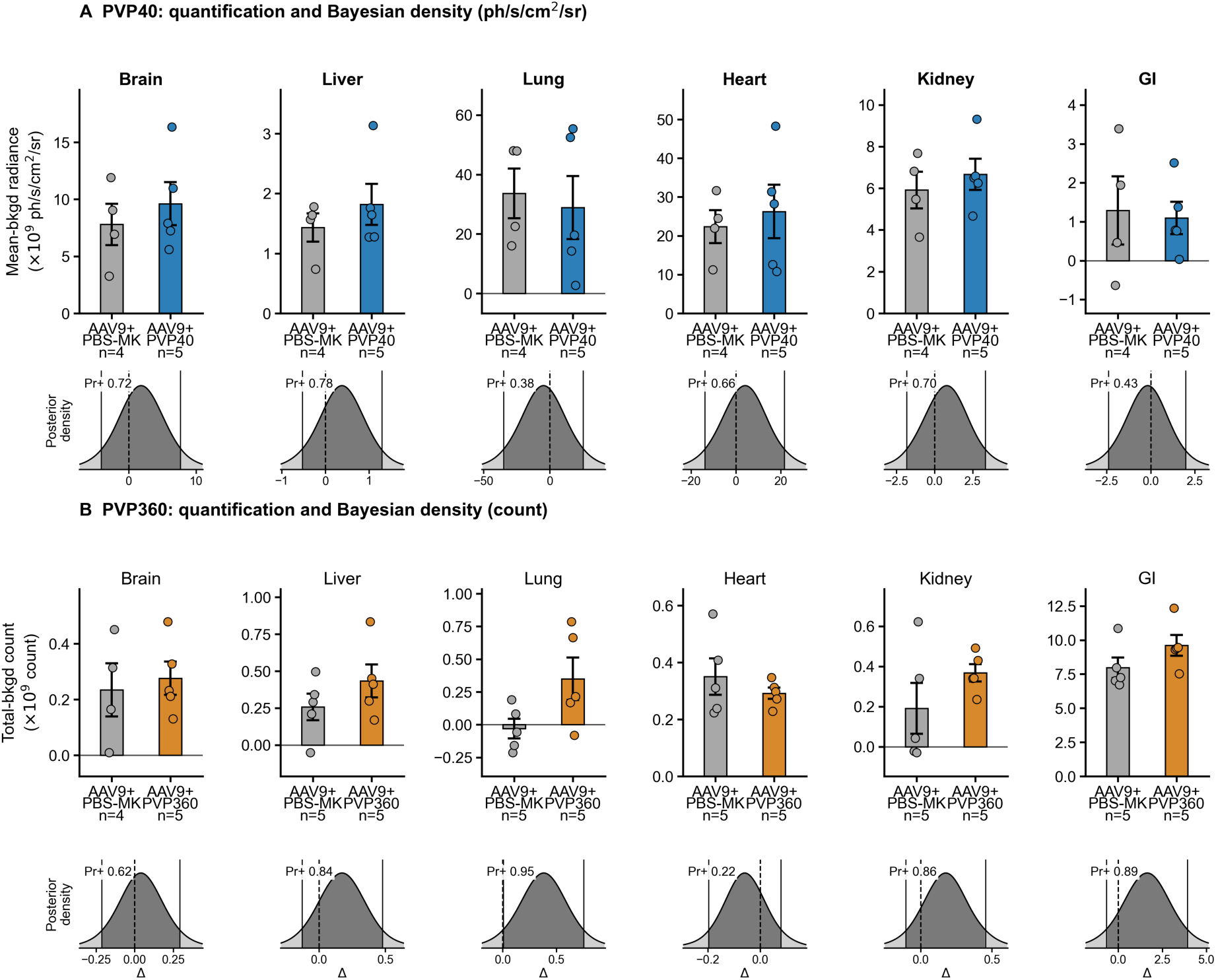
*Ex vivo* organ reporter-signal quantification after intravenous delivery with Bayesian posterior summaries. (A) PVP40-cohort data for six organs, shown as background-subtracted radiance and posterior density plots for treatment-minus-AAV9+PBS-MK mean differences (control n = 4; PVP40 n = 5). (B) PVP360-cohort data for six organs, shown as background-subtracted total counts and posterior density plots (n = 5/group except brain control n = 4). Background-subtracted values, including negative values, were retained on the source scale. PVP40 and PVP360 were acquired in separate cohorts with separate AAV9+PBS-MK controls and different quantitative metrics and were not compared directly across panels. PVP360 showed an isolated positive posterior signal in lung, whereas the remaining organ comparisons did not provide consistent evidence of enhancement.

**Supplementary Figure 6.**
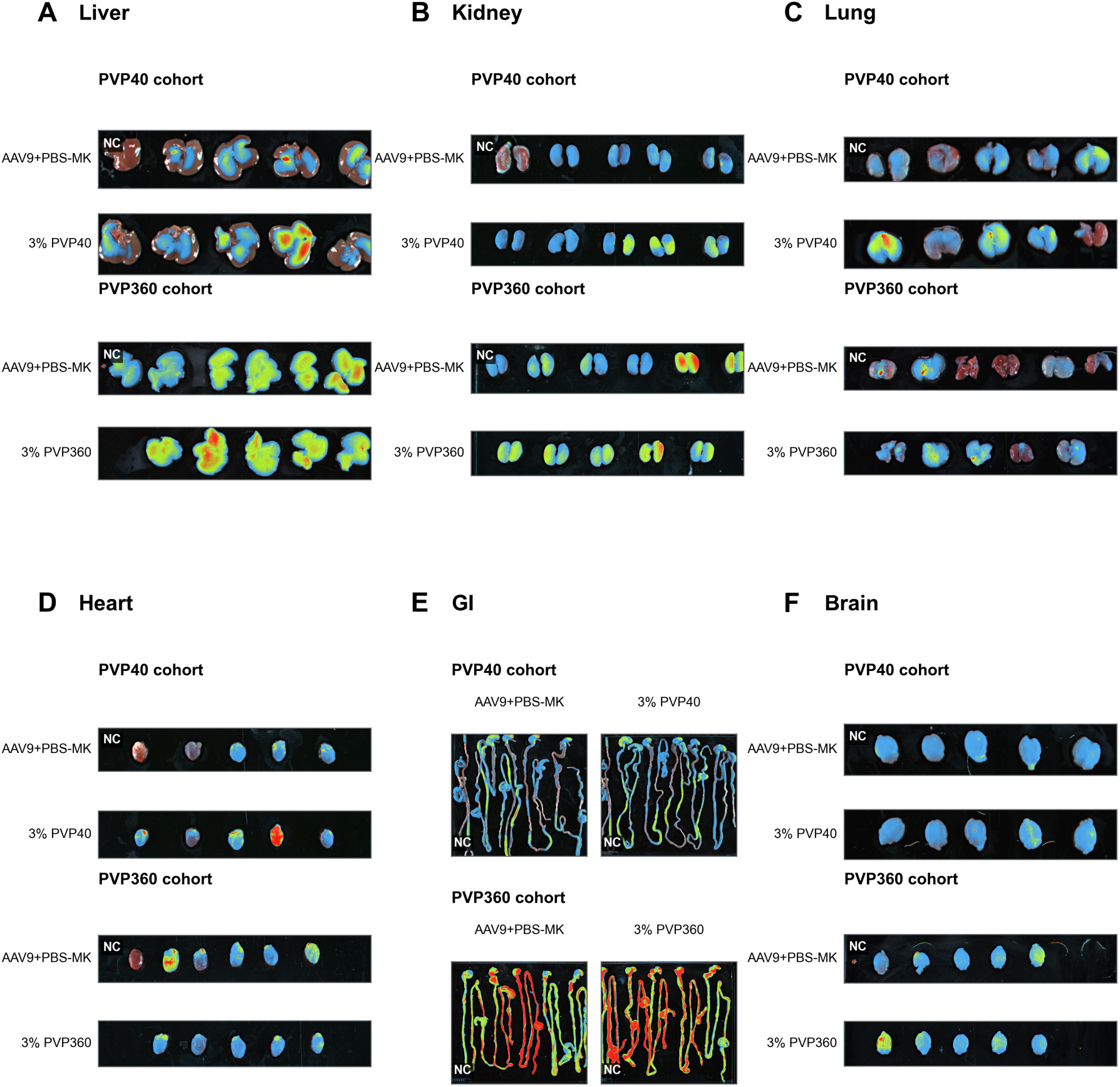
Representative *ex vivo* organ reporter images after intravenous delivery. Panels show (A) liver, (B) kidney, (C) lung, (D) heart, (E) gastrointestinal tract, and (F) brain. Within each panel, images from the PVP40 cohort show the AAV9+PBS-MK control and 3% PVP40 groups, and images from the separate PVP360 cohort show the AAV9+PBS-MK control and 3% PVP360 groups. NC denotes an organ from an uninjected negative-control mouse included at the left of the corresponding specimen row or GI field. Quantification and Bayesian posterior summaries are shown in Supplementary Figure 5.

**Supplementary Figure 7.**
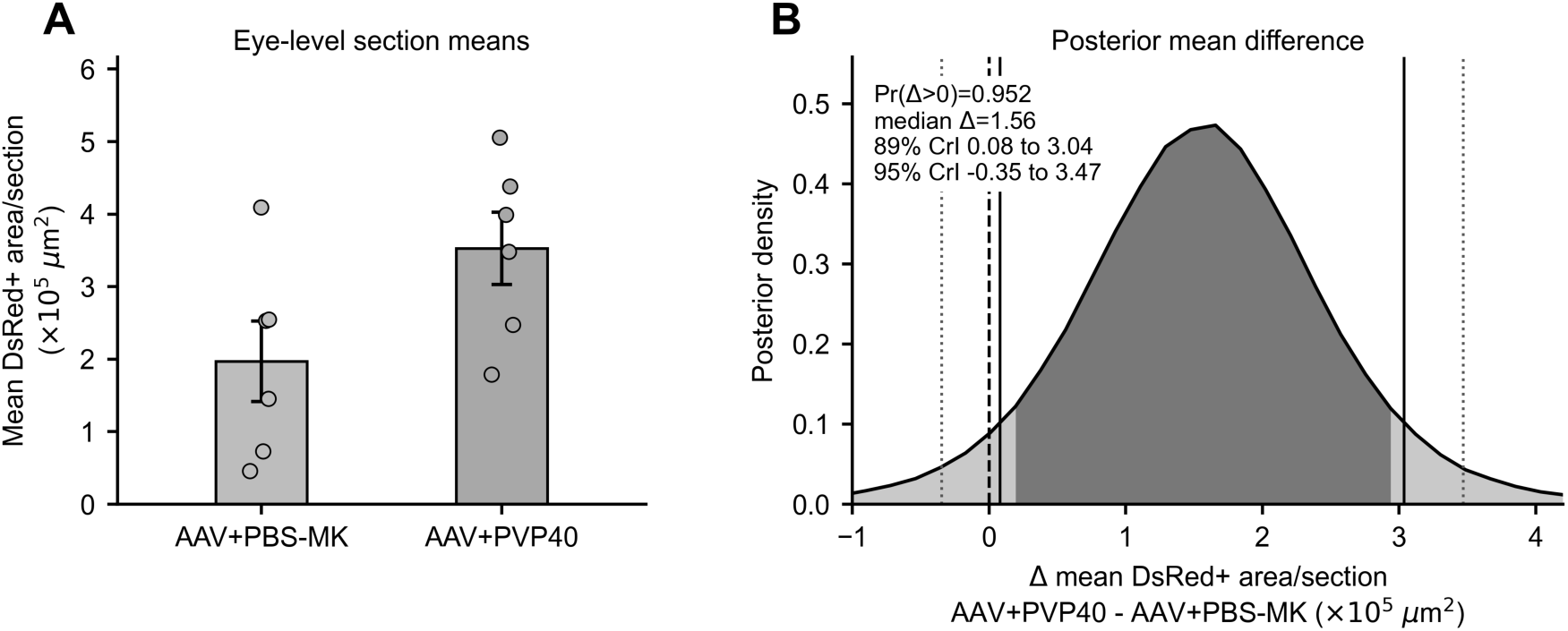
Bayesian sensitivity analysis for local subretinal delivery. (A) The n = 6 eyes/group mean DsRed-positive area per retinal section values used in Figure 4B are shown in grayscale. (B) Posterior distribution of the AAV+PVP40-minus-AAV+PBS-MK group mean difference under the reference-prior normal model described in the Methods. Bayesian analysis favored a positive PVP40 mean difference (Pr[Δ > 0] = 0.952; median Δ = 1.56 × 10^5^ µm^2^; central 89% credible interval, 0.08 to 3.04 × 10^5^ µm^2^; central 95% credible interval, -0.35 to 3.47 × 10^5^ µm^2^).

**Supplementary Figure 8.**
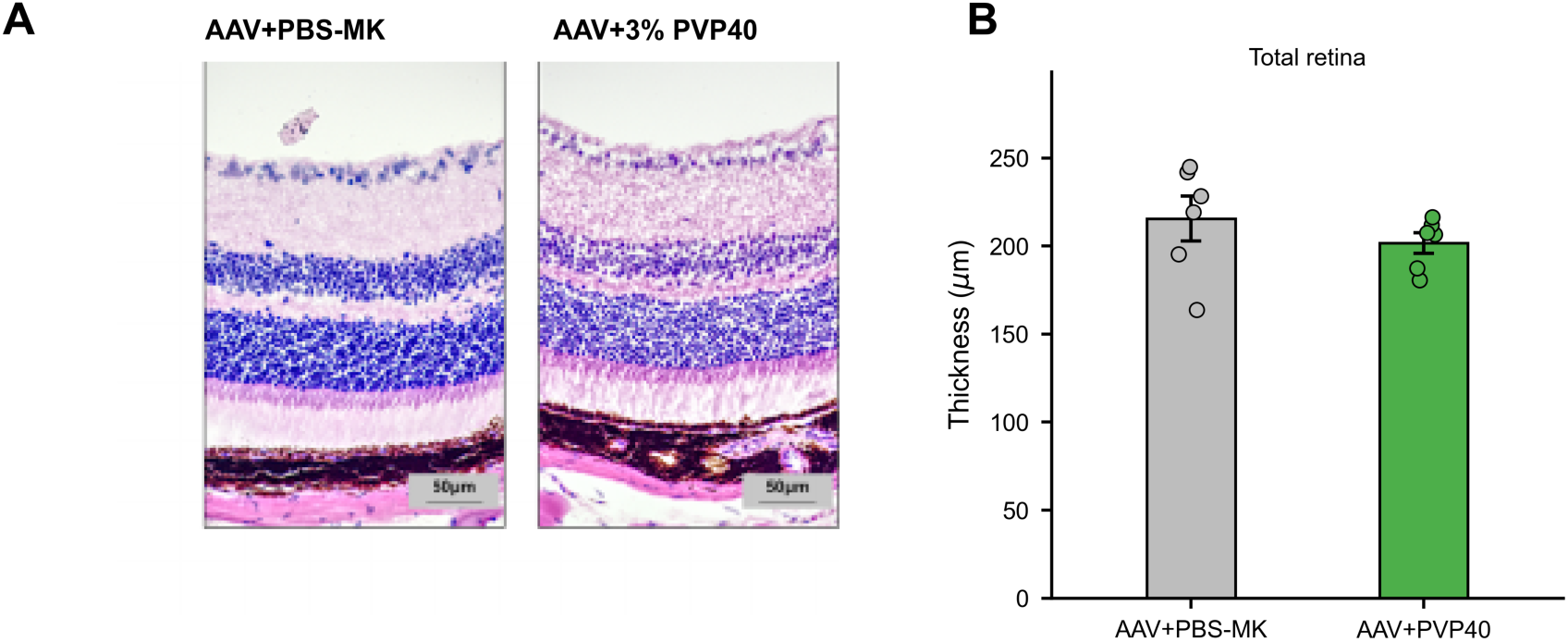
Retinal histology two weeks after subretinal delivery. (A) Representative H&E retinal sections from AAV+PBS-MK vehicle and AAV+3% PVP40 groups. (B) Total retinal thickness from n = 6 eyes/group; three retinal positions were averaged for each eye.

## Supplementary Tables

**Supplementary Table S1.**
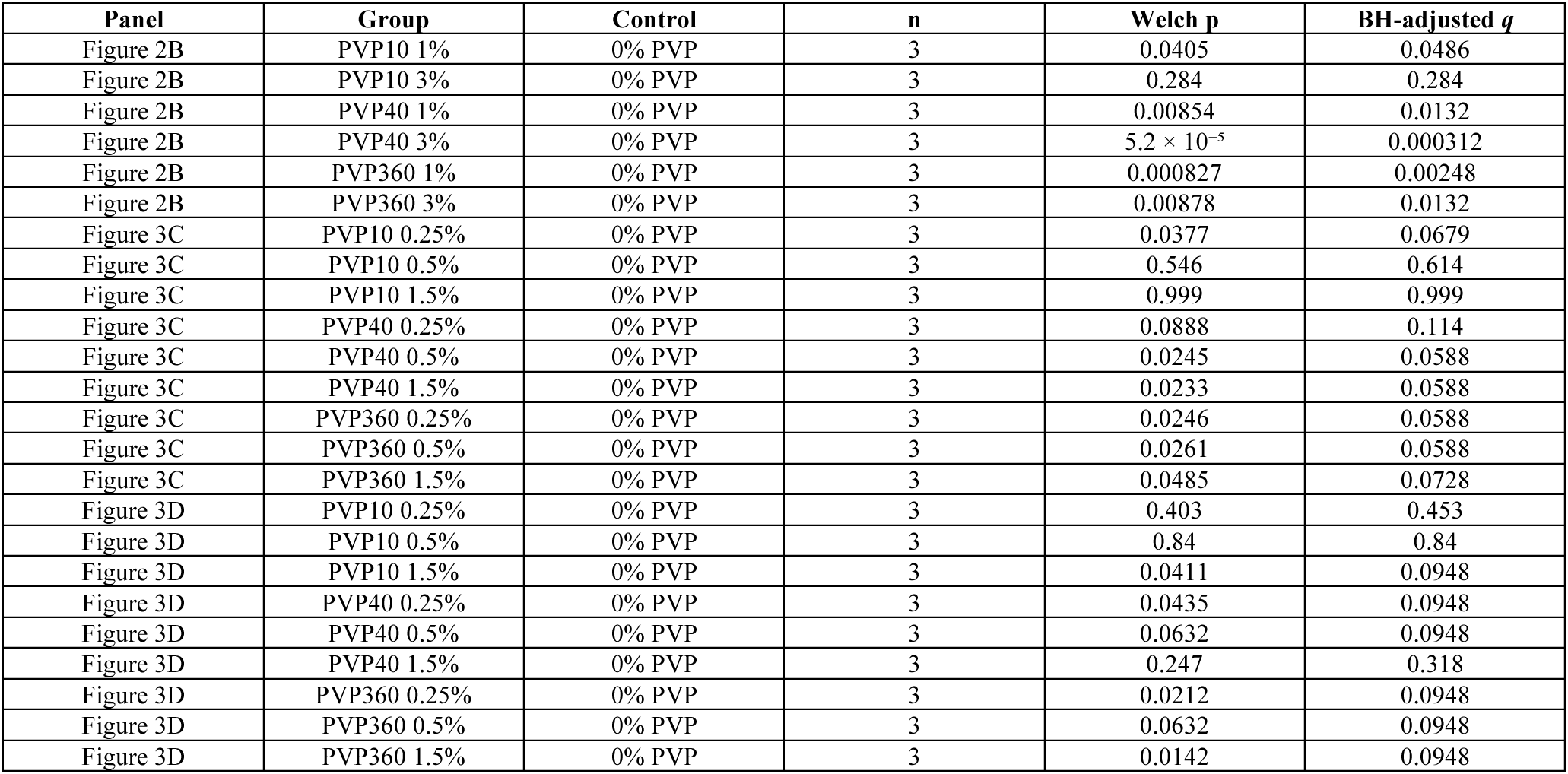
Welch’s t-tests and Benjamini-Hochberg-adjusted *q* values for Figure 2 and Figure 3 screening panels. Figure 2B asterisks correspond to the BH-adjusted *q* values versus the 0% PVP control; Figure 3 values are tabulated without figure asterisks. n indicates the number of replicate samples per group.

